# Maternal alcohol consumption during pregnancy and offspring epigenome-wide DNA methylation: findings from six general population-based birth cohorts

**DOI:** 10.1101/167791

**Authors:** Gemma C Sharp, Ryan Arathimos, Sarah E Reese, Christian M Page, Janine Felix, Leanne K Küpers, Sheryl L Rifas-Shiman, Chunyu Liu, The Cohorts for Heart and Aging Research in Genomic Epidemiology plus (CHARGE +) methylation alcohol working group, Kimberley Burrows, Shanshan Zhao, Maria C Magnus, Liesbeth Duijts, Eva Corpeleijn, Dawn L DeMeo, Augusto Litonjua, Andrea Baccarelli, Marie-France Hivert, Emily Oken, Harold Snieder, Vincent Jaddoe, Wenche Nystad, Stephanie J London, Caroline L Relton, Luisa Zuccolo

**Affiliations:** MRC Integrative Epidemiology Unit, University of Bristol, Bristol, UK; School of Social and Community Medicine, University of Bristol, Bristol, UK; School of Oral and Dental Sciences, University of Bristol, Bristol, UK; Division of Intramural Research, National Institute of Environmental Health Sciences, National Institutes of Health, Department of Health and Human Services, Research Triangle Park, North Carolina, USA; Department of Non-Communicable diseases, Division for Mental and Physical Health, Norwegian Institute of Public Health, Oslo, Norway; Oslo Centre for Biostatistics and Epidemiology, Oslo University Hospital, Oslo, Norway; The Generation R Study Group, Erasmus MC, University Medical Centre Rotterdam, Rotterdam, the Netherlands; Department of Epidemiology, Erasmus MC, University Medical Centre Rotterdam, Rotterdam, the Netherlands; Department of Pediatrics, Erasmus MC, University Medical Centre Rotterdam, Rotterdam, the Netherlands; Department of Epidemiology, University of Groningen, University Medical Center Groningen, Groningen, the Netherlands; Department of Population Medicine, Harvard Medical School, Harvard Pilgrim Health Care Institute, Boston, MA, USA; The Framingham Heart Study, Framingham, MA, USA.; The Population Sciences Branch, Division of Intramural Research, National Heart, Lung, and Blood Institute, Bethesda, MD, USA.; Boston University School of Public Health, 715 Albany St, Boston, MA, USA.; Department of Pediatrics, Division of Respiratory Medicine and Allergology, Erasmus MC, University Medical Centre Rotterdam, Rotterdam, the Netherlands; Department of Pediatrics, Division of Neonatology, Erasmus MC, University Medical Centre Rotterdam, Rotterdam, the Netherlands; Channing Division of Network Medicine, Brigham and Women’s Hospital, Harvard Medical School, Boston, MA USA; Laboratory of Precision Environmental Biosciences, Columbia University Mailman School of Public Health, New York, NY, USA; Diabetes Unit, Massachusetts General Hospital, Boston, MA, USA

## Abstract

Some evidence suggests that light-to-moderate alcohol consumption during pregnancy is associated with adverse outcomes in the offspring, but the precise biological mechanisms underlying such associations are currently unknown. Epigenetic modifications have been suggested as one potential explanation.

Within the Pregnancy and Childhood Epigenetics (PACE) consortium, we performed meta-analysis to combine information from six population-based birth cohort studies to investigate DNA methylation at over 450,000 sites in the cord blood of newborns differentially exposed to alcohol *in utero*. We were primarily interested in the effects of sustained consumption throughout pregnancy (data available for five cohorts, 3,075 mother-child pairs), which represents a prolonged prenatal exposure to alcohol, but we also explored binge-drinking and timing-specific exposures. In addition to looking for differential methylation at individual CpG sites, we also used two different methods, Comb-P and DMRcate, to identify differentially methylated regions (DMRs).

We found no strong evidence of association between any of our alcohol exposure measures and DNA methylation at any individual CpG site. Using Comb-P, we identified 19 DMRs in the offspring of mothers who drank throughout pregnancy compared to the offspring of mothers who gave up drinking at the start of pregnancy, but these were not validated using DMRcate.

In this multi-cohort study of the general population we found no evidence that maternal alcohol consumption during pregnancy is associated with offspring cord blood DNA methylation, which is in stark contrast to the multiple, strong associations that previous studies have found for maternal smoking. However, it is possible that a combination of a larger sample size, higher doses, different timings of exposure and a more global assessment of genomic DNA methylation might show evidence of association.

## Introduction

It is well known that heavy alcohol consumption during pregnancy can cause Fetal Alcohol Spectrum Disorders (FASD), a spectrum of disorders characterized by a continuum of structural and neurodevelopmental abnormalities, with Fetal Alcohol Syndrome (FAS) at the more severe end of the spectrum [1–3]. The severity of FASD appears to depend largely on the timing, dose and frequency of exposure to alcohol, with heavy exposure in the latter half of the first trimester being associated with the most severe effect [4,5]. However, in the general population most pregnant women do not drink at the doses required to cause FASD. For example, in population-based studies from Ireland, the UK, Australia and New Zealand, around 70% (range 67% to 77%) of women who reported drinking in the first trimester consumed seven units or fewer per week, which is considered light-to-moderate consumption. In the second trimester, nearly all women who drank (range 99% to 100%) consumed seven units or fewer per week [6]. Evidence of an effect of light-to-moderate levels of prenatal alcohol exposure is sparse and inconsistent. Although the majority of systematic reviews and studies published after these reviews have not found convincing evidence of association between light-to-moderate drinking and adverse offspring health and neurodevelopment [7–14], a recent comprehensive review of prospective studies found suggestive evidence of an association between mothers consuming up to four UK units of alcohol per week and babies born small-for-gestational age or preterm [15]. Furthermore, results from quasi-experimental study designs, which are more robust to the presence of confounding by parental socio-economic factors, have shown some evidence of effect of (mostly light-to-moderate) maternal alcohol consumption on offspring cognition and behaviour [16–18]. The inconsistency of findings may be explained by the failure to adequately control for certain confounding factors, such as socioeconomic position, diet and ethnicity [19], which affect offspring outcomes both prenatally and postnatally, and could therefore bias any potentially small effect of light-to-moderate drinking in pregnancy [20]. Suggestions of harm from these later studies, together with findings from animal experiments, have prompted the UK Chief Medical Officer to recently revise the guidelines for alcohol drinking in pregnancy to recommend abstention, based on the precautionary principle [21].

Whether there is a causal association between light-to-moderate drinking in pregnancy and children’s health outcomes is an important question. Identifying a possible biological pathway showing effects at birth would be a first step towards providing an answer. Currently, precise biological mechanisms underlying potential adverse effects of prenatal alcohol exposure are unknown. However, epigenetic modifications have been suggested as one such potential mediator, with some evidence that this is the case for prenatal exposure to smoking [22–24].

Animal studies suggest that alcohol exposure affects DNA methylation levels both globally, through its antagonistic effect on methyl donors such as folate [25,26], and in a gene-specific fashion [27,28]. Mouse pups exposed to alcohol during the highly epigenetically-sensitive intrauterine period show dose- and timing-specific epigenetic effects, including DNA methylation effects that correlate with long-lasting changes in gene expression and could potentially drive offspring adverse outcomes [29].

Similar experiments are impossible in humans for obvious ethical reasons. However, there is some evidence that treatment with a low physiologically relevant dose of ethanol induces genome-wide changes in DNA methylation in human embryonic stem cells [30]. In addition, a recent observational study of 110 children with FASD and 96 controls found genome-wide differences in buccal epithelial cell DNA methylation [31]. It is still unknown whether light-to-moderate prenatal alcohol exposure is associated with differential DNA methylation in human offspring.

In this study, we meta-analysed epigenome-wide association study (EWAS) summary statistics from six population-based cohort studies within the Pregnancy and Childhood Epigenetics (PACE) Consortium to investigate DNA methylation profiles in the cord blood of newborns differentially exposed to alcohol *in utero*. We also compared these associations with those recently found in studies of 1) differential buccal cell DNA methylation in children with FASD compared to controls [31], and 2) differential whole blood DNA methylation in adults in the general population who drink light-to-moderately compared to adults who do not drink [32].

## Methods

### Participating cohorts

A total of six independent cohorts from four countries participated in this study, all were members of the PACE Consortium. Detailed methods for each cohort are provided in the Supplementary Material (File S1). All cohorts had data on maternal alcohol consumption before and/or during pregnancy and DNA methylation data as measured using the Illumina Infinium HumanMethylation450k BeadChip array [33]. In alphabetical order, these cohorts were: The Avon Longitudinal Study of Parents and Children (ALSPAC) [34–37] from the UK, Groningen Expert Center for Kids with Obesity (GECKO) [38] and Generation R (GENR) [39,40] from the Netherlands, two independent datasets from the Norwegian Mother and Child Cohort Study (MoBa1, MoBa2) [41,42], and Project Viva (Viva) from the USA [43].

### Maternal alcohol consumption (exposure)

Cohorts assessed maternal alcohol consumption before and during pregnancy via questionnaires completed by the mothers during pregnancy. We were primarily interested in the effects of sustained consumption throughout pregnancy, which represents a longer prenatal exposure to alcohol, potentially interfering with all stages of embryonic and fetal development. Therefore, our main exposure of interest was a binary variable comparing offspring of mothers who drank both before pregnancy and in the second and/or third trimester of pregnancy to offspring of mothers who consumed alcohol before pregnancy but not during the second and/or third trimester of pregnancy. Using this definition, we hoped to compare offspring of mothers who continued to drink alcohol after finding out they were pregnant to offspring of mothers who stopped drinking. From previous research we know that women who drink in the second trimester tend to do so at light-to-moderate levels [6].

Cohorts ran secondary models assessing binge drinking during pregnancy and timing-specific alcohol consumption: before pregnancy, during the first trimester and during the second and/or third trimester. Binge drinking was defined as four, five or six (depending on the cohort) or more glasses per occasion at least once at any time point in pregnancy compared to consuming alcohol before pregnancy and drinking in moderation (i.e. no binge drinking) during pregnancy. Alcohol consumption before pregnancy, during the first trimester and during the second and/or third trimester were all defined using four categories of exposure: 1) no drinking, 2) less than one glass per week, 3) one to six glasses per week, 4) seven or more glasses per week.

### Covariates

All models were adjusted for maternal age (years), maternal education (variable defined by each individual cohort) and maternal smoking status (the preferred categorisation was into three groups: no smoking in pregnancy, stopped smoking in early pregnancy, smoking throughout pregnancy. A binary categorisation of any versus no smoking was also acceptable). All cohorts also adjusted for technical covariates either by including a batch variable (for example chip ID) as a model covariate or by generating and adjusting for surrogate variables. All models were run with and without adjustment for cell counts, which were estimated using the Houseman method [44]. The analyses were completed before a cord blood reference panel was widely available, so cohorts used an adult whole blood reference [45] to estimate the proportion of B-cells, CD8+ T-cells, CD4+ T-cells, granulocytes, NK-cells and monocytes in each sample.

### Methylation measurements (outcome)

DNA from cord blood underwent bisulfite conversion using the EZ-96 DNA methylation kit (Zymo Research Corporation, Irvine, USA). DNA methylation was measured using the Illumina Infinium HumanMethylation450k BeadChip assay at Illumina or in cohort-specific laboratories. Each cohort conducted its own quality control and normalisation of methylation data, as detailed in the Supplementary Material (File S1). In all analyses, cohorts used normalised, untransformed beta-values. As a consortium, we have found that extreme outliers in methylation data, likely caused by technical error or rare genetic variants, can have a large influence on results. Therefore, potential outliers were removed. Such outliers were defined using the Tukey method [46], in which an outlier is any value less than the lower quartile minus three times the interquartile range, or more than the upper quartile plus three times the interquartile range. This method is appropriate as it is not dependent on distributional assumptions of the data.

### Cohort-specific statistical analysis

Each cohort performed independent epigenome-wide association studies (EWAS) according to a common, pre-specified analysis plan. Models were run using multiple robust linear regression (rlm in the MASS R package [47]) in an attempt to control for potential heteroscedasticity in the methylation data. Alcohol consumption was modelled as the exposure and cord blood DNA methylation was the outcome, with adjustment for covariates (and estimated cell counts).

### Meta-analysis

We performed fixed-effects meta-analysis weighted by the inverse of the variance with METAL [48]. We then excluded control probes (N=65) and probes mapped to the X (N=11,232) or Y (N=416) chromosomes. This left a total of 473,864 probes. Multiple testing was accounted for by controlling the false discovery rate (FDR) at 5% using the method by Benjamini and Hochberg [49]. Probes were annotated according to hg19 using the IlluminaHumanMethylation450kanno.ilmn12.hg19 R package [50].

### Sensitivity analyses

For the top 500 sites with the smallest P-values in our main model (sustained drinking), we repeated the meta-analysis using a random effects model, to allow for potential differences in effect sizes between cohorts. We additionally assessed inter-study heterogeneity and influence of individual cohorts by observing forest plots and heterogeneity statistics, as well as conducting a “leave-one-out” analysis using the metafor R package [51]. We checked consistency between models by comparing effect estimates and top hits to those of our primary model. We also compared top hits to a list of probes suggested to give spurious readings due to cross-hybridisation or genomic features such as nearby SNPs [52]. When a cord blood reference became available [53], we ran a sensitivity analysis in ALSPAC adjusting for estimated proportions of B-cells, CD8+ T-cells, CD4+ T-cells, granulocytes, NK-cells, monocytes and nucleated red blood cells.

### Comparison to associations between FASD and buccal epithelial DNA methylation in children

We performed a look-up in our results of the top CpGs and differentially methylated regions (FDR-adjusted P-value<0.05, n CpGs = 658) reported in a study by Portales-Casamar *et al.* [31], which analysed epigenome-wide buccal epithelial cell DNA methylation in children with FASD compared to controls. We compared direction of effect and p-values across the two studies.

### Comparison to associations between light-to-moderate drinking and whole blood DNA methylation in adults

We performed a second look-up in our results of the top CpGs (FDR-adjusted P-value<0.05) from a study by Liu *et al.* [32], which meta-analysed associations between alcohol consumption and epigenome-wide whole blood DNA methylation in adults (CHARGE consortium). In order to harmonise with our models, we restricted the look-up to the top CpGs from the previous study’s models that assessed light drinking (3 CpGs) and moderate drinking (24 CpGs), versus no drinking (results supplied by the study authors). As with our first look-up, we compared direction of effect and p-values across the two studies.

### Differentially methylated regions analysis

Adjacent probes on the 450k array are often highly correlated and differentially methylated regions (DMRs) may be more biologically important than individual CpGs. Therefore, we used two methods, Comb-P [54] and DMRcate [55], to identify DMRs in our meta-analysed single-CpG EWAS results. Comb-P identifies genomic regions enriched for low P-values, corrects for auto-correlation with neighbouring CpGs within 1000 bp using the Stouffer-Liptak method, then adjusts for multiple testing using the Sidak correction. DMRcate generates two smoothed estimates for each chromosome: one weighted by F-statistics (calculated from the meta-analysis results as (beta/standard error)^2^) and one not, for null comparison. The two estimates are compared via a Satterthwaite approximation and P-values are calculated and adjusted for multiple testing using the FDR method. Regions are defined from groups of significant probes (FDR<0.05) where the distance to the next consecutive probe is less than 1000 bp. A regional P-value is calculated using the Stouffer method. As a sensitivity analysis, we repeated the Comb-P and DMRcate analyses using a 500 bp (rather than 1000 bp) window to define neighbouring CpGs.

### Blood/Brain comparison

To assess whether identified associations between prenatal alcohol exposure and cord blood methylation are likely to represent associations in a more biologically-relevant tissue (the brain), we performed a look-up of CpG sites in a database of correlations between blood and brain methylation (http://epigenetics.iop.kcl.ac.uk/bloodbrain/). Methods used to derive this database are described elsewhere [56], but briefly, the authors quantified DNA methylation in matched DNA samples from whole blood and four brain regions (prefrontal cortex, entorhinal cortex, superior temporal gyrus and cerebellum) from 122 individuals.

## Results

### Study characteristics

Of the six participating cohorts, Project Viva and GECKO could not take part in all meta-analyses due to insufficient data/number of exposed individuals. Project Viva were therefore only included in the analysis of drinking before pregnancy compared to abstaining. GECKO were included in the primary analysis assessing sustained alcohol consumption, and the secondary analysis assessing binge drinking. The other cohorts (ALSPAC, Generation R, MoBa1 and MoBa2) were included in all analyses.

Five cohorts (ALSPAC, GECKO, Generation R, MoBa1, MoBa2) had the necessary data to take part in our primary analysis of sustained alcohol consumption in pregnancy. The meta-analysis included 3,075 mother-child pairs, of which 1,147 (37.3%) mothers consumed alcohol both before and throughout pregnancy and the remaining 1,928 mothers consumed alcohol before pregnancy/during the first trimester but not during the second and/or third trimester. Table 1 summarizes the characteristics of each cohort. In all investigated cohorts, women who drank throughout pregnancy were, on average, older and had a higher level of education than women who stopped. In MoBa2 and GECKO women who drank throughout pregnancy were less likely to smoke throughout pregnancy compared to women who stopped drinking, but the opposite was true for the other three studies.

**Table 1.** Summaries of cohorts and meta-analysis results for the sustained maternal alcohol consumption model among 3,075 mother-newborn pairs across five studies.

|  | Exposed* |  |  |  | Unexposed* |  |  |  | EWAS results<br>unadjusted for cell<br>counts |  | EWAS results<br>adjusted for cell<br>counts |  |
| --- | --- | --- | --- | --- | --- | --- | --- | --- | --- | --- | --- | --- |
| Cohort | N | Mean<br>maternal<br>age (SD) | N any<br>smoking in<br>pregnancy | N high<br>education<br>level | N | Mean<br>maternal<br>age (SD) | N any<br>smoking in<br>pregnancy | N high<br>education<br>level | Lambda | N CpGs<br>with FDR-corrected<br>P<0.05 | Lambda | N CpGs<br>with FDR-corrected<br>P<0.05 |
| ALSPAC | 540 | 30.2 (4.1) | 69 (13%) | 131 (24%) | 235 | 28.6 (4.7) | 25 (11%) | 36 (15%) | 1.07 | 0 | 1.08 | 0 |
| GECKO | 70 | 31.5<br>(4.21) | 30 (43%) | 35 (50%) | 66 | 30.4<br>(4.05) | 32 (48%) | 19 (29%) | 0.90 | 2 | 0.98 | 2 |
| Generation R | 440 | 32.5 (3.7) | 111 (25%) | 349 (79%) | 283 | 30.6 (4.3) | 59 (21%) | 152 (54%) | 1.27 | 0 | 1.14 | 0 |
| MoBa 1 | 242 | 31.4 (3.9) | 75 (31%) | 178 (74%) | 632 | 29.3 (4.3) | 190 (30%) | 416 (66%) | 0.77 | 0 | 0.65 | 0 |
| MoBa 2 | 157 | 31.8 (4.2) | 39 (25%) | 115 (73%) | 414 | 29.3 (4.4) | 112 (71%) | 238 (58%) | 2.20 | 0 | 1.63 | 0 |
| *Total* | *1147* | *31.5* | *324* | *808* | *1928* | *29.6* | *418* | *861* | *1.04 †* | *0 †* | *1.17 †* | *0 †* |

\*Mothers in the 'exposed' group drank both before pregnancy and in the second and/or third trimester. Mothers in the 'unexposed' group drank before pregnancy but not after the first trimester.
† Based on results from a meta-analysis using a fixed-effects model

### Primary models: sustained maternal alcohol consumption during pregnancy

For both the cell-adjusted and cell-unadjusted models, effect sizes were moderate: for top CpGs with P-value<1*10^-3^, estimates ranged from a 4% decrease to 2% increase in average methylation level in the exposed compared to the unexposed group, with a median absolute estimate of 0.4% (File S2, Table S1 and S2). No CpG sites survived correction for multiple testing with an FDR-adjusted P-value<0.05 (Figure 1). There was little evidence of inflation, as assessed by the lambda value (Table 1). Results for all sites with a P-value<1*10^-3^ are presented in File S2 Table S1 and S2. Of the 622 and 797 CpGs with P-value<1*10^-3^ in the cell-unadjusted and cell-adjusted models, respectively, 500 had a P-value<1*10^-3^ according to both models. Adjusting for estimated cell counts appeared to have little effect: at the 500 top sites, the median percentage difference in effect sizes before and after adjustment was 3.4% and only 26/500 sites changed by 10% or more. At exactly half of the 500 top sites, adjusting for cell counts reduced the effect size towards the null. At the remaining 250 sites the effect size increased after cell-adjustment. Adjusting for cell counts increased the standard error at 357/500 sites.

**Figure 1.**
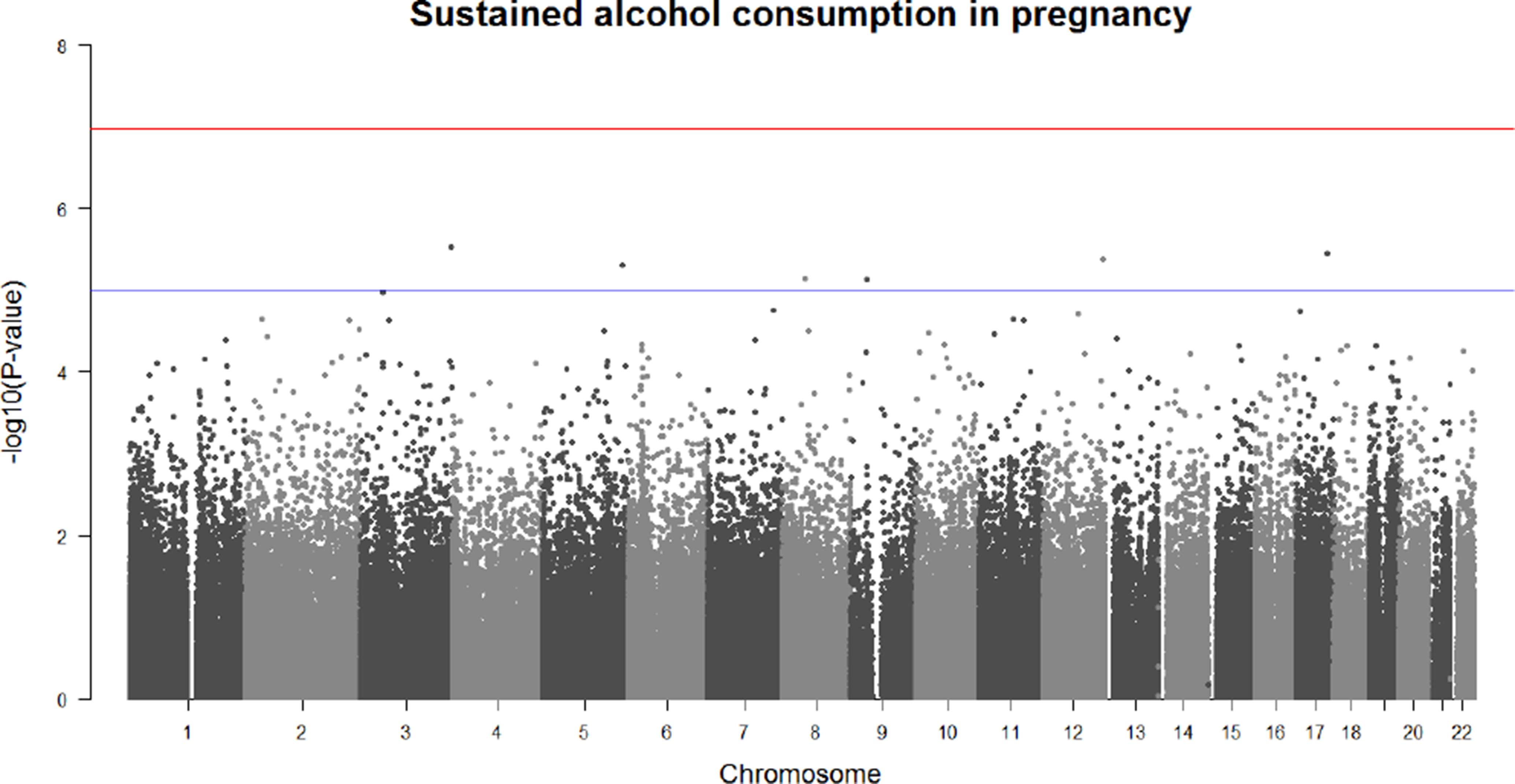
Manhattan plot of associations between sustained maternal alcohol consumption and offspring cord blood DNA methylation (without adjustment for cell counts). The top (red) line shows the -log10 P-value threshold for Bonferroni significance. The bottom (blue) line shows the -log10 P-value threshold for FDR significance.

### Secondary models

With and without adjustment for cell counts, no individual CpG sites were associated with drinking before pregnancy or during the first trimester after FDR correction for multiple testing. One CpG (cg12509712 near *ARSG*) was associated with binge drinking, but only after adjustment for cell counts. In addition, one CpG (cg20334115 near *PYCR2*) was associated with drinking in the second and/or third trimester, but only before adjustment for cell counts. We did not consider these individual sites further due to the lack of consistency between the cell-adjusted and cell-unadjusted models. Results for all sites with a P-value<1*10^-3^ are presented in File S2 Table S3-S10.

### Sensitivity analyses

There was evidence of heterogeneity at a minority of the top 500 sites associated with sustained maternal alcohol consumption: 100/500 sites had a heterogeneity P-value<0.05; I^2^ was >40 at 36/500. After running a random effects meta-analysis at the top 500, the coefficients for 88/500 changed >10% compared to coefficients generated using the fixed effects meta-analysis. Forest plots and results of a leave-one-out analysis (File S3) suggested that no single cohort had a disproportionately large influence on the meta-analysis results consistently over all 500 sites.

Of the top 500 sites, 164 were on a published list of possibly problematic probes [52] (File S2 Table S1 and S2). Although these sites may be more likely to contain outliers, cohorts removed outlier values prior to EWAS and used a regression model that is designed to be robust to outliers in the outcome variable.

The direction of effect in our primary model (sustained maternal alcohol consumption during pregnancy) was mostly consistent with results from our other models (Figure 2). As is expected given their similarity, results from the model assessing maternal alcohol consumption in the second/third trimester were particularly consistent with results from the primary model (Spearman’s correlation coefficient for regression coefficients at all probes: 0.76 for the cell-adjusted models and 0.74 for the cell-unadjusted models). Results for all sites with a P-value<1*10-3 are presented in File S2 Table S1-S10 and lambdas and number of hits per cohort are provided in File S2 Table S11.

**Figure 2.**
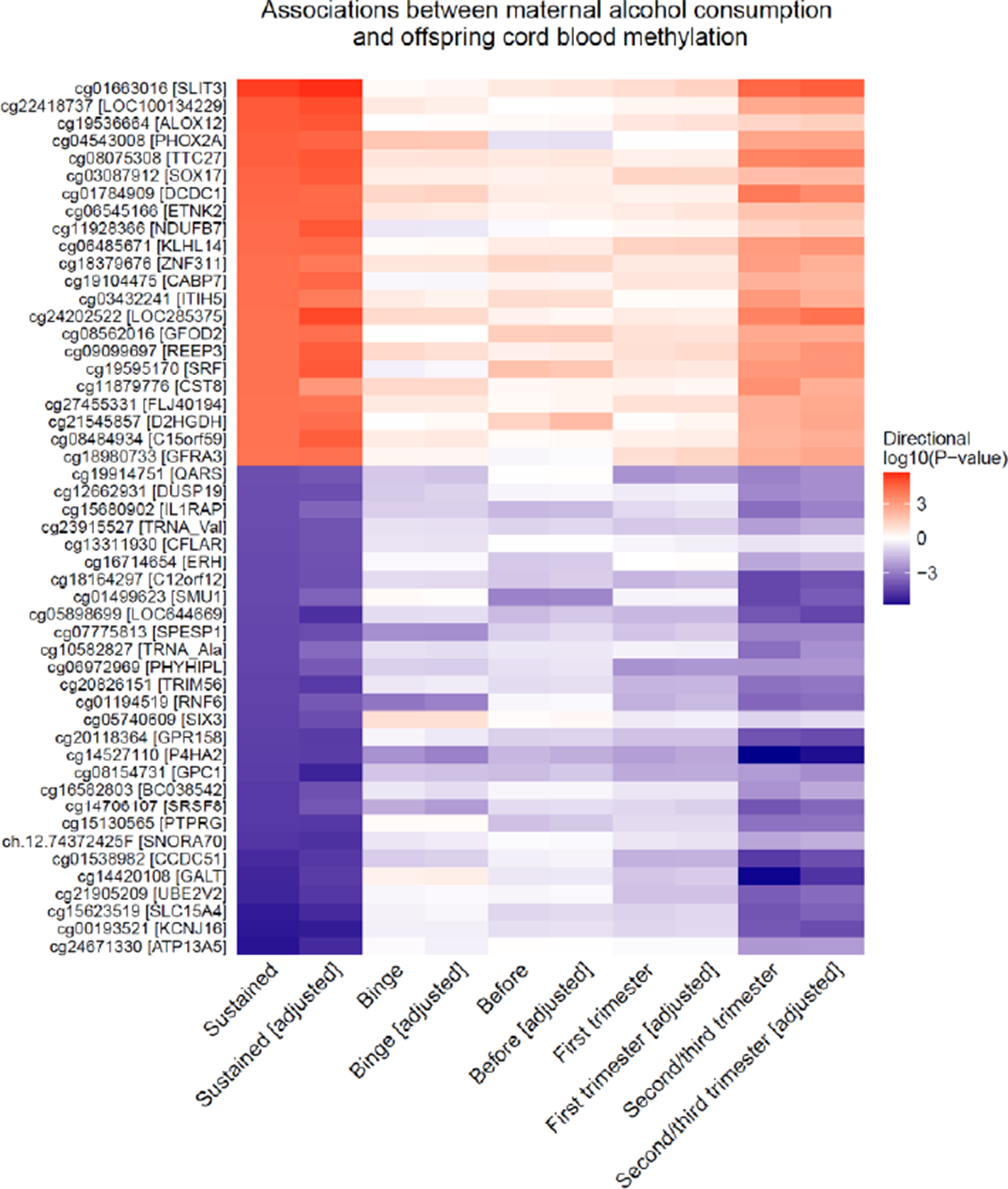
A heatmap to illustrate the direction and strength of association between all investigated alcohol exposures and offspring cord blood DNA methylation. Plotted CpGs are the top 50 CpGs with the smallest P-values in the sustained alcohol consumption single-site EWAS (without adjustment for estimated cell proportions). “adjusted” denotes models that were adjusted for estimated cell proportions.

In ALSPAC, adjusting for cell counts estimated using the adult whole blood reference panel provided similar EWAS results to those obtained when adjusting for cell counts estimated using a cord blood reference panel: at the top 500 sites from the primary meta-analysis, the median percentage change in coefficients in ALSPAC was 7% (IQR 3% to 14%), with 317/500 sites changing less than 10%. Of the top 500 sites, 219 had a crude P-value < 0.05 in ALSPAC when using the adult reference, and 197/219 (90%) also had a crude P-value <0.05 when using the cord blood reference.

### Comparison to associations between FASD and buccal epithelial DNA methylation in children

Of 658 CpG sites at which DNA methylation in buccal cells was associated with FASD according to Portales-Casamar *et al*., 288 (44%) had the same direction of association (regardless of P-value) in our study of maternal sustained drinking, but none survived correction for multiple testing at 658 sites (FDR-adjusted P-value<0.05). Of 542 CpG sites within 101 FASD-associated DMRs (identified using DMRcate), 215 (40%) had the same direction of association in our study of maternal sustained drinking, but none survived correction for multiple testing at 542 sites (FDR-adjusted P-value<0.05). Full look-up results for all PACE models are provided in File S2 Table S12.

### Comparison to associations between light/moderate drinking and whole blood DNA methylation in adults

Of 24 CpGs associated (FDR-adjusted P<0.05) with adult moderate drinking in the study by Liu *et al.* [32], one was associated with maternal sustained drinking in our PACE analysis after correction for multiple testing at 24 sites (FDR-adjusted P<0.05). At this site, the direction of the effect of maternal drinking on methylation was inverse compared to the effect of own drinking: cg19909613; closest gene TTC35; PACE effect 0.0075 P-value 1.7*10^-3^; Liu *et al.* effect -0.014 P-value 9.2*10^-7^. In the cell-adjusted model, this site did not survive FDR correction. None of the three CpG sites that were associated with light drinking according to Liu *et al.* were associated with maternal sustained drinking in the PACE study. Full look-up results for all PACE models are provided in File S2 Table S13.

### Differentially methylated region analysis

Using Comb-P to conduct a region-based analysis based on spatial correlation of p-values, we identified 30 and 32 DMRs in the cell-unadjusted and cell-adjusted maternal sustained drinking models, respectively (Sidak-corrected P-value <0.05). Nineteen regions were differentially methylated according to both models (Table 2; results for all models shown in File S2 Table S14). However, we found no DMRs when we conducted the region-based analysis using DMRcate. Defining DMRs using a 500bp (as opposed to 1000bp) window did not change our results using either Comb-P or DMRcate.

**Table 2.** Regions identified using the Comb-P method as differentially methylated in association with sustained maternal alcohol consumption.

| Differentially-methylated region (DMR) | CpGs in DMR | Closest gene | Sidak P-value (unadjusted for cell counts) | Sidak P-value (adjusted for cell counts) |
| --- | --- | --- | --- | --- |
| Chr1: 152161237-152162026 | 7 | *HRNR* | 1.0E-13 | 6.0E-11 |
| Chr16: 1583810-1584517 | 8 | *IFT140* | 8.7E-13 | 5.9E-10 |
| Chr6: 49681178-49681775 | 9 | *CRISP2* | 9.5E-11 | 7.6E-08 |
| Chr5: 110062343-110062838 | 7 | *TMEM232* | 2.1E-10 | 2.0E-07 |
| Chr17: 6899085-6899759 | 12 | *ALOX12* | 1.3E-09 | 9.3E-07 |
| Chr12: 31271783-31272120 | 4 | *DKFZp434C0631* | 8.8E-09 | 1.2E-05 |
| Chr19: 57741988-57742445 | 10 | *AURKC* | 1.2E-08 | 1.2E-05 |
| Chr8: 43131260-43131657 | 5 | *POTEA* | 3.8E-07 | 4.5E-04 |
| Chr5: 8457548-8458090 | 6 | *LOC100505738* | 5.5E-07 | 4.8E-04 |
| Chr13: 76334583-76334867 | 4 | *LMO7* | 5.5E-07 | 4.8E-04 |
| Chr5: 42953543-42953625 | 3 | *AK056817* | 9.4E-07 | 1.6E-03 |
| Chr2: 118594304-118594650 | 3 | *DDX18* | 1.2E-06 | 7.1E-03 |
| Chr3: 145879277-145879711 | 7 | *PLOD2* | 1.5E-06 | 2.1E-03 |
| Chr17: 35423405-35423817 | 4 | *AATF* | 2.4E-06 | 2.6E-03 |
| Chr18: 77659572-77659696 | 2 | *KCNG2* | 4.1E-06 | 4.7E-03 |
| Chr15: 69222988-69223369 | 3 | *SPESP1* | 4.6E-06 | 1.7E-02 |
| Chr2: 103236861-103237269 | 2 | *SLC9A2* | 6.1E-06 | 7.6E-03 |
| Chr6: 31846769-31847029 | 10 | *SLC44A4* | 6.4E-06 | 7.4E-03 |
| Chr4: 3516534-3516759 | 4 | *LRPAP1* | 9.2E-06 | 1.7E-02 |

### Blood/brain comparison

CpGs within the 19 sustained drinking DMRs identified using Comb-P tended to show strong correlations between blood and brain methylation. For example, at the top CpG with the smallest EWAS P-value within the top DMR with the smallest SIDAK-corrected P-value (cg26320663, DMR Chr1:152161237-152162026 near *HRNR*) correlation coefficients between blood and brain ranged from 0.61 in the cerebellum and entorhhinal cortex to 0.66 in the pre-frontal cortex (P-values ranging 1.8*10^-8^ to 1.5*10^‒10^). All results are presented in File S2 Table S15.

## Discussion

We combined data across five pregnancy cohorts to evaluate associations between maternal alcohol consumption during pregnancy and genome-wide DNA methylation in cord blood of offspring. Although we did not find any consistent evidence of association between any category of prenatal alcohol exposure, defined in terms of dose and timing, and single-site methylation, we found some evidence that 19 larger regions of the genome were differentially methylated in association with sustained maternal alcohol consumption throughout pregnancy (i.e. when comparing mothers who drank throughout pregnancy to those who stopped at the beginning or after the first trimester). However, optimal methods for regional analyses are currently a matter of debate and we were not able to validate this finding using a different region-based method, so we conclude that we have not found any strong evidence for an association.

The only previously published study on methylation effects of prenatal alcohol exposure was reported by Portales-Casamar *et al*., who compared buccal cells from children with and without FASD in a small sample with just over 100 cases [31]. At the CpGs that Portales-Casamar *et al.* identified as associated with FASD, less than half showed the same direction of association with sustained maternal alcohol consumption in our study, and none survived correction for multiple testing. This is perhaps not surprising given the stark methodological differences between the two studies: Firstly, DNA methylation is strongly tissue-specific [57], so DNA from buccal cells (Portales-Casamar *et al*.) and cord blood (our study) are perhaps unlikely to show the same general methylation patterns. Secondly, DNA methylation is strongly influenced by age [58] and could be affected by many factors in the postnatal environment that are associated with prenatal alcohol exposure (such as maternal education and childhood adversity) – both of these factors are important to consider when interpreting results from our study, where methylation was measured at birth, compared to the study by Portales-Casamar *et al*., where participants were around 11-years-old. Thirdly, the prospective cohort design of the studies included in our meta-analysis meant that we were able to adjust for measured confounders and participants were sampled from the same populations. Portales-Casamar *et al*.’s case-control study of 11-year-olds is more open to confounding (especially by the postnatal environment, as mentioned above). In particular, the cases and controls differed with respect to important sociodemographic characteristics (namely, ethnicity and being raised by adoptive/foster parents) that have previously been associated with variation in DNA methylation [59–61]. Fourthly, the FASD case-control comparison likely covers a larger range of exposure (i.e. cases were exposed to much higher intensity of exposure in *utero*; controls were likely to have been exposed to less alcohol) than our study of differential prenatal exposure to alcohol in the general population. This is supported by previous population-based studies showing that women who drink during pregnancy tend to do so at light-to-moderate levels, especially those that continue to drink after pregnancy detection [6]. Fifthly, this difference in level of alcohol exposure in both studies means there are different confounding structures, for example, in Portales-Casamar *et al.* higher exposure was associated with lower socioeconomic status, whereas, in our study, higher exposure was associated with higher maternal education in all investigated cohorts.

We were interested in whether exposure to alcohol prenatally is associated with DNA methylation at the same CpG sites as exposure to alcohol during adulthood. Therefore, we performed a look-up of results from a study recently published by Liu *et al.*, which found some evidence of associations between DNA methylation and varying levels of alcohol consumption in an adult population [32]. At the CpGs that were associated with light or moderate drinking in Liu et *al.*, we did not find any strong evidence of an association with prenatal exposure. This lack of overlap might be explained by maternal alcohol consumption affecting fetal DNA methylation differently than adult methylation. It could also be related to differences in population age, definitions, range and duration of exposure, and potential differences in accuracy of self-report in non-pregnant and pregnant adults leading to differences in measurement error.

Although we did not find strong evidence for an association between alcohol consumption during pregnancy and genome-wide DNA methylation in offspring cord blood in the general population, this does not exclude the possibility that such an association does exist in a different tissue. For example, brain tissue may be a more appropriate tissue to study because some of the strongest evidence from observational studies suggests an association between prenatal alcohol exposure and impaired neurodevelopmental outcomes [62]. There are obvious ethical issues that preclude collection of brain tissue in population-based studies, however, there are reasons to believe that DNA methylation in blood may be a good surrogate for DNA methylation in brain at some sites. Strong correlations in blood and brain DNA methylation have been found at some CpGs [56,63], including at our top DMR from the Comb-P analysis. Although we were not able to replicate our Comb-P DMRs using a different method (DMRcate), the high correlation between some of these regions in blood and brain mean they may serve as interesting candidate genes for further studies of the role of DNA methylation as a mediator of associations between prenatal alcohol exposure and offspring neurodevelopmental outcomes. Alternatively, the high correlations between blood and brain methylation at these sites might represent a genetic influence on methylation that is also associated with maternal alcohol consumption. Regardless of causality, methylation at these regions might be a useful biomarker for prenatal alcohol exposure [64], thereby serving as a more objective measure than self-report. This possibility would have to be validated and tested in independent datasets.

Strengths of our study include the use of data from six well-characterised and established cohorts. The prospective data available from these cohorts has allowed us to investigate the timing and strength of exposure, including binge-drinking and exposure by trimester, and to minimise measurement error and recall bias. Previous studies from the PACE consortium have used a similar methodology to prescribe cohort-specific analyses and to meta-analyse results from several cohorts. These studies identified many, seemingly robust, associations [22,65]. This suggests that the lack of associations identified in our study is not likely due to poor methodology or data.

Failure to observe associations between maternal alcohol consumption and cord blood DNA methylation could be due to lack of statistical power, particularly if we hypothesise a dose-response association such that low levels of exposure correspond to small methylation differences. As most pregnant women in our cohorts did not drink excessive amounts of alcohol, a large sample size would be required to detect a small epigenetic effect. A further potential limitation of our study is that there was some inter-cohort heterogeneity, perhaps caused by differences in confounding structures (we note that in some studies women who drank throughout pregnancy were more likely to be smokers, while in others the opposite was true), definition and range of alcohol exposure, and/or different methods of normalizing DNA methylation data. Encouragingly though, our meta-analysis results were not substantially different when we used a random-effects model compared to a fixed-effects model. Furthermore, forest plots and results of a leave-one-out analysis suggested our meta-analysis results were not strongly influenced by differences between studies. Measurement error could also reduce statistical power and this may be a particular problem for studies of maternal alcohol consumption: pregnant women may under-report behaviours that are widely thought to be harmful for their baby [66]. Cellular heterogeneity in cord blood samples is a further issue [67] that may introduce error in our measure of DNA methylation. Blood samples are highly heterogeneous and although we have attempted to adjust for cellular heterogeneity by including estimated cell proportions in our EWAS models, no suitable cord blood reference was widely available at the time of analysis, so these estimates were based on an adult blood reference panel. When a cord blood reference became available [53], a sensitivity analysis in ALSPAC did not reveal sizable differences between EWAS results adjusted for cell proportions estimated using either the cord or adult reference panel. However, we recognise that there may be a residual influence of cellular heterogeneity that could be biasing the results in either direction. Another factor that could be biasing results is circulating folate levels, which we could not formally evaluate because not all cohorts had the required data. However, there are two reasons why folate is likely to play at most a marginal role in our findings. Firstly, we found low inter-study heterogeneity for most of our CpG sites – if there was a prominent interaction between alcohol (a folate antagonist) and folate (a methyl donor), then we might expect more heterogeneity due to country- and timing-specific differences in folate intake. Secondly, our results were largely null – confounding by folic acid supplementation, which is indirectly associated with alcohol consumption [14,68], is unlikely to be exaggerating estimates of the association between maternal alcohol and offspring methylation. We also consider that some of our findings may be affected by selection bias: women who are actively trying to get pregnant may abstain from drinking alcohol, but these women were excluded from our analyses of sustained and binge drinking. However, these women were not excluded from our analyses of other alcohol exposures (before pregnancy, first trimester, second and third trimester) and our findings were consistently null across all alcohol exposures. A final potential limitation is that the Illumina HumanMethylation450k array covers only 1.7% of all CpG sites in the human genome, which may not cover potential regions where maternal alcohol consumption is most strongly associated with offspring methylation. The development of sequence-based approaches methods with better genomic coverage will help overcome this limitation in future studies.

If an association between maternal alcohol consumption and cord blood DNA methylation does exist (for example, in larger samples and/or different populations with a greater range of alcohol exposure), then future studies should explore whether that association is causal or confounded by genetics or shared mother-child environmental factors. For example, paternal alcohol consumption could be employed as a ‘negative control’ that will share the same confounding structure as maternal alcohol consumption but cannot plausibly affect offspring DNA methylation through a direct causal intrauterine mechanism [69]. Techniques such as two-step Mendelian randomization [70] could also be applied to explore 1) the causal effect of prenatal alcohol exposure on DNA methylation and 2) the causal effect of DNA methylation on offspring outcomes. Even if the association is not causal, newborn blood DNA methylation might capture both genetic and environmental influences of maternal alcohol consumption, which could be useful, both clinically and in research, as a biomarker of exposure and/or a useful predictor of offspring outcomes.

## Conclusion

In this multi-cohort study, we found no evidence that maternal alcohol consumption during pregnancy is associated with offspring cord blood DNA methylation in the general population. However, it is possible that a combination of higher doses and different timings of exposure, as well as a more global assessment of genomic DNA methylation, might show evidence of association. We therefore recommend caution when interpreting the present null findings and encourage further investigations.

